# Combined Disruption of Multiple Cytokine Signaling Pathways Enables One-Step Anterograde Tracing with Vesicular Stomatitis Virus

**DOI:** 10.64898/2026.05.22.727255

**Authors:** Xiang Ma, Constance L. Cepko

**Affiliations:** Department of Genetics, Harvard Medical School, Boston, MA 02115; Howard Hughes Medical Institute, Chevy Chase, MD 20815

**Author notes:** **Corresponding Author:** Constance L. Cepko.

## Abstract

Defining the direct postsynaptic targets of selected neuronal populations remains a major challenge for neural circuit mapping. Vesicular stomatitis virus (VSV) spreads efficiently in the anterograde direction, but replication-competent VSV undergoes multistep spread and therefore cannot distinguish direct from indirect downstream targets. Here, we developed a glycoprotein-deleted VSV(VSVdG)-based strategy for one-step anterograde tracing using AAV-mediated trans-complementation with several adaptations. In this system, VSVdG was engineered to encode Cre, allowing a Cre-dependent AAV to express VSV-G only after VSVdG infected the same cells, thereby limiting VSV-G expression to a short time window. To reduce VSV-M-mediated cytotoxicity, we introduced the M33A/M51R double-mutant VSV-Md variant. Using the basal ganglia circuit as a model, we found that efficient one-step VSVdG spread required disruption of cytokine-mediated antiviral responses triggered by AAV transduction and VSV infection. This was achieved by loss of type I interferon signaling in IFNAR1-knockout mice, together with suppression of additional antiviral responses either through AAV-mediated expression of the CVS-N2c rabies virus phosphoprotein or administration of cytokine-blocking antibody cocktail. These adaptations enabled VSVdG spread from dorsal striatum to expected downstream targets in mice of both sexes. However, the downstream labeled cells included both neurons and glia, revealing an important limitation for interpreting this approach as strictly neuron-to-neuron monosynaptic anterograde spread. Overall, rather than providing a ready-to-use tracing tool, this study presents a VSVdG strategy for one-step anterograde circuit tracing, and defines viral toxicity, innate immunity, and cell-type specificity constraints that must be addressed to develop an efficient and specific monosynaptic anterograde viral tracer.

**Significance Statement:** Mapping direct downstream targets of defined neuronal populations is essential for understanding neural circuit function, but reliable monosynaptic anterograde viral tracers remain limited. We developed a VSVdG-based strategy that uses AAV-mediated trans-complementation to restrict VSV-G expression to starter cells in a short time window and incorporates VSV-M variant to reduce toxicity. In the mouse basal ganglia, this system enabled one-step spread from dorsal striatum to expected output regions when innate antiviral barriers were suppressed. Our results identify type I interferon and additional type I/type II interferon-independent cytokine signaling as major restrictions of VSVdG spread. This study establishes proof of principle for VSV-based one-step anterograde tracing while defining viral toxicity, innate immunity, and cell-type specificity constraints in need of further improvement.

## Introduction

Mapping neural circuit connectivity is essential for understanding the principles underlying normal neural function and neurological disorders. Viral tracers are powerful tools for defining synaptic connections, and monosynaptic tracers are especially valuable because they identify direct inputs to, or outputs from, genetically defined starter neurons. Glycoprotein-deleted rabies virus (RABVdG) has been widely used for monosynaptic retrograde tracing of direct neuronal inputs (Callaway & Luo 2015, Reardon et al 2016, Wickersham et al 2007b). In contrast, viral tools for monosynaptic anterograde tracing of direct neuronal outputs remain limited, although several studies have attempted to address this challenge (Fischer et al 2023, Li et al 2021, Rivera et al 2025, Tsai et al 2022, Xiong et al 2022, Zeng et al 2017, Zingg et al 2017).

Replication-competent recombinant vesicular stomatitis virus (VSV) undergoes robust anterograde spread through the nervous system, labeling multiple brain regions across diverse organisms (Beier et al 2011, Drokhlyansky et al 2017, Kler et al 2021, Lin et al 2020, Mundell et al 2015). However, because replication-competent VSV undergoes multistep spread, it cannot distinguish direct from indirect downstream targets. Developing a VSV-based system for monosynaptic anterograde tracing would therefore provide an important complement to existing monosynaptic anterograde and retrograde tracers.

VSV and RABV are both enveloped, nonsegmented, negative-sense RNA viruses in the family *Rhabdoviridae*, although they belong to distinct genera, *Vesiculovirus* and *Lyssavirus*, respectively (Walker et al 2018). Both viruses have the genomic organization 3’-N-P-M-G-L-5’ and in both viruses the G gene encodes the only transmembrane envelope glycoprotein. The glycoprotein is not required for replication of the viral genome but is essential for receptor recognition and for delivery of the viral genome into infected cells (Lichty et al 2004, Schnell et al 2010). Thus, a conceptually straightforward strategy for monosynaptic anterograde tracing with VSV is to adapt the trans-complementation approach used for RABVdG-based retrograde tracing, in which glycoprotein-deleted virus spreads only from starter cells that supply G in trans by AAV.

However, we made repeated attempts for complementation of VSVdG by an AAV encoding VSV-G, and found that this approach was insufficient for monosynaptic anterograde tracing *in vivo*. This failure likely reflects fundamental biological differences between VSV and RABV. First, high-level constitutive expression of VSV-G is toxic in mammalian cells (Burns et al 1993, Chen et al 1996), in contrast to RABV-G, which has been used to generate stable producer cell lines for RABVdG amplification (Osakada & Callaway 2013, Wickersham et al 2010), Second, RABV evades innate immunity mainly through RABV-P and RABV-M, which antagonize specific antiviral signaling pathways (Kiflu 2024, Schnell et al 2010, Scott & Nel 2016). In contrast, VSV relies on VSV-M– mediated global host shutoff as a major immune-evasion strategy (Ahmed & Lyles 1998, Petersen et al 2000, von Kobbe et al 2000). These activities of VSV-G and VSV-M could compromise starter-cell viability and interfere with AAV expression.

VSV-M variants with impaired host-shutoff activity provide a potential strategy to reduce cytotoxicity, but this attenuation also increases VSV sensitivity to innate antiviral responses. Because both AAV and VSV can induce type I interferons and pro-inflammatory cytokines that have antiviral function (Georgel et al 2007, Hosel et al 2012, Jiang et al 2005, Shao et al 2018, Shi et al 2011, Zhu et al 2009), efficient anterograde spread of VSVdG encoding attenuated VSV-M variants likely requires suppression of cytokine-mediated immune signaling. Here, we sought to identify barriers that restrict VSVdG-based anterograde tracing *in vivo* and to test adaptations to overcome them. We used the basal ganglia pathway as a model circuit because striatal projection neurons send well-defined outputs to the globus pallidus and substantia nigra (Nelson & Kreitzer 2014), and previous work from our laboratory demonstrated robust VSV-based multistep anterograde tracing in this circuit (Beier et al 2011, Drokhlyansky et al 2017, Mundell et al 2015). Using this system, we evaluated how VSV-G expression timing, VSV-M attenuation, cytokine signaling, and microglia influence VSVdG-mediated anterograde spread.

## Materials and Methods

### Animals

All animal procedures were approved by the Institutional Animal Care and Use Committee (IACUC) of Harvard University. Mice were housed under standard conditions on a 12 h light/12 h dark cycle with food and water available *ad libitum*. Both male and female mice, 8–33 weeks of age, were used.

The following mouse lines were used in this study: C57BL/6J (000664, The Jackson Laboratory); IFNAR1 knockout mice (032045-JAX in Figure 1 and 028288 in all other experiments, The Jackson Laboratory); IFNGR1 knockout mice (003288, The Jackson Laboratory); IFNAR1/IFNGR1 double-knockout (DKO) mice (generated by crossing 032045-JAX and 003288 for Figure 1 and 029098 for all other experiments, The Jackson Laboratory); Myd88 knockout mice (009088, The Jackson Laboratory); Trif^Lps2^ mice (005037, The Jackson Laboratory); and MAVS knockout mice (008634, The Jackson Laboratory). The number of animals used in each experiment is indicated in figure legends.

**Figure 1.**
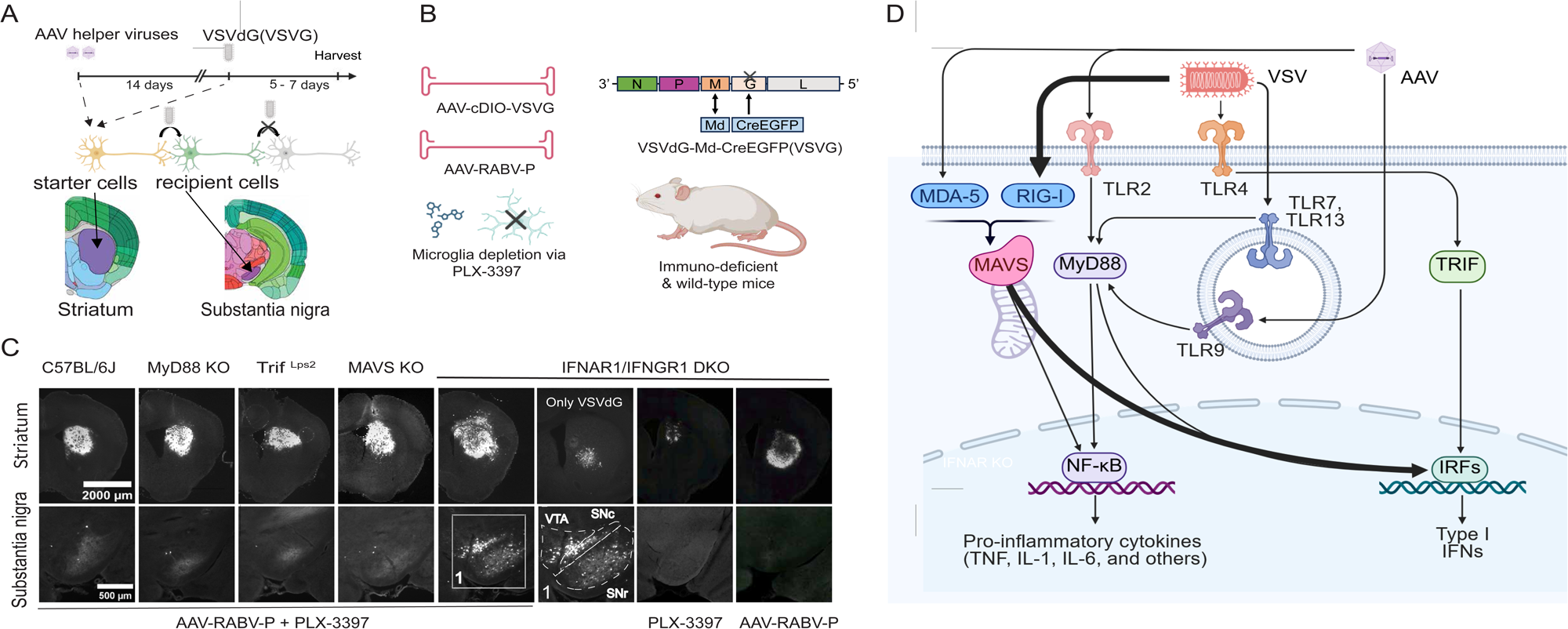
Requirements for one-step anterograde spread with VSVdG-Md-CreEGFP(VSV-G) (A) Experimental design (B) The AAVs, VSV, microglia-depletion drug, and mouse lines used in this experiment. (C) Representative coronal sections of the dorsal striatum, substantia nigra, and ventral tegmental area (VTA) showing EGFP labeling after VSV infection under the indicated conditions. In IFNAR1/IFNGR1 double-knockout mice, four experimental conditions were examined: AAV-RABV-N2cP without microglia depletion, microglia depletion without AAV-RABV-N2cP, AAV-RABV-N2cP combined with microglia depletion, and a VSV-only control that received VSVdG-Md-CreEGFP(VSV-G) without either AAVs or microglia depletion. The first three experimental groups also received AAV-VSV-G. In C57BL/6, Myd88-knockout, Trif-deficient, and MAVS-knockout mice, only the combined AAV-VSV-G, AAV-RABV-N2cP, and microglia-depletion condition was tested. n = 2 each. (D) Diagram showing that AAV and VSV are sensed by multiple innate immune pathways, including Toll-like receptors (TLRs) and RIG-I-like receptors (RLRs), which trigger production of type I interferons and pro-inflammatory cytokines through the adaptor proteins MyD88, MAVS, and TRIF.

### Cell lines

293T cells (ATCC® CRL-3216™), 293T–TVA800 cells (gift from John Young, Salk Institute), and BHK-21 cells (EH1011, Kerafast) were cultured in Dulbecco’s modified Eagle’s medium (11995065, Thermo Fisher Scientific) supplemented with 10% heat-inactivated fetal bovine serum (FBS, 10437028, Thermo Fisher Scientific) and 1 U/ml penicillin-streptomycin (15140-163, Invitrogen) at 37 °C and 5% CO_2_.

### Plasmids

All plasmids were constructed by Gibson assembly. The vesicular stomatitis virus vectors used in this study were based on the Indiana serotype (Beier et al 2011, Drokhlyansky et al 2017, Krause & Cepko 2024). A glycoprotein-deleted VSV genomic plasmid encoding CreEGFP and carrying the M33A/M51R double mutation in VSV-M was generated and is referred to here as VSVdG-Md-CreEGFP. A replication-competent VSV genomic plasmid with the organization 3′-TagBFP-N-P-Mq-G-H2BEGFP-L-5′, in which Mq denotes the M33A/M51R/V221F/S226R quadruple mutation in VSV-M, was also generated and is referred to here as VSV-Mq-H2BEGFP (Hoffmann et al 2010). Plasmids used for VSV rescue encoded VSV N, P, L, M, G, and T7 polymerase. All rescue plasmids, with the exception of the VSV genomic plasmids, were gifts from Christopher L. Parks (International AIDS Vaccine Initiative, IAVI). pCMV-loxp-TVA-mCherry-ARE-loxp-EnvA/VSVG contains an AU-rich element (ARE) sequence, UUAUUUAUUGAUCCUUAU UUAUU, that destabilizes TVA-mCherry mRNA (Zubiaga et al 1995). The EnvA/VSV-G chimera was generated by fusing the extracellular and transmembrane domains of EnvA to the intracellular tail of VSV-G. The EnvA sequence was derived from Addgene plasmid #32206. In the plasmids used to generate AAV, the VSV-G sequence used was mouse codon-optimized and synthesized by IDT. The N2cP sequence was derived from Addgene plasmid #100808, the B19P sequence from Addgene plasmid #32631, and the TVA-mCherry sequence from Addgene plasmid #48332.

### Virus production and preparation

VSVdG and replication-competent VSV were rescued using a modified version of a previously described protocol (Witko et al 2006). Briefly, 293T cells were grown to 100% confluence in a 6-cm dish, dissociated, and seeded into poly-D-lysine-coated 6-well plates at a 1:3 split ratio. Cells were maintained in DMEM supplemented with 10% FBS and 1% penicillin-streptomycin at 37 °C with 5% CO₂. The following day, cells were washed with PBS and transfected in each well with plasmids encoding VSV-N (400 ng), VSV-P (300 ng), VSV-L (100 ng), VSV-M (250 ng), VSV-G (250 ng), T7 polymerase (10 μg), and the VSV genomic plasmid (5 μg). Transfection was performed using linear polyethylenimine (PEI, 25 kDa; 23966; Polysciences) at a PEI:DNA ratio of 2:1 in 500 μl DMEM per well. After 4–6 h, the medium was replaced with 5 ml fresh DMEM to remove PEI, and cells were maintained at 32 °C with 3% CO₂. The following day, cells were heat-shocked at 43 °C with 3% CO₂ for 3 h and then returned to 32 °C with 3% CO₂. This 3-h heat shock at 43 °C was repeated daily until a strong EGFP signal was observed.

VSVdG-Md-CreEGFP(VSV-G) was amplified from rescued virus stocks. Briefly, BHK-21 cells grown to 100% confluence in a 10-cm dish were dissociated and seeded into a new 10-cm dish at a 1:15 split ratio. Cells were maintained in DMEM supplemented with 10% FBS and 1% penicillin-streptomycin at 37 °C with 5% CO₂. On the following day, the medium was replaced with 12 ml DMEM alone, and cells were transfected with 3 μg pCMV-VSV-G plasmid per dish. Transfection was performed using linear polyethylenimine (PEI, 25 kDa) at a PEI:DNA ratio of 4:1 to 6:1 in 3 ml DMEM per dish. Twenty-four hours later, the medium was replaced with 2 ml DMEM, and 50 μl of supernatant from the rescue well was added to each dish. After 1 h, 13 ml DMEM. Viral supernatant was harvested 2 days later.

VSVdG-Md-CreEGFP(EnvA) was amplified from VSVdG-Md-CreEGFP(VSV-G), except that the amplification plasmid was changed from pCMV-VSV-G to pCMV-loxp-TVA-mCherry-ARE-loxp-EnvA. Briefly, BHK-21 cells in 10-cm dishes were transfected with 6 μg pCMV-loxp-TVA-mCherry-ARE-loxp-EnvA per dish. Twenty-four hours later, the medium was replaced with 2 ml DMEM, and cells were infected with VSVdG-Md-CreEGFP(VSV-G) at a multiplicity of infection (MOI) of 0.01. After 1 h, 13 ml DMEM was added to each dish. Viral supernatant was harvested 3-4 days post-infection, when viral spread had reached the entire dish.

For VSVdG-Md-CreEGFP(VSV-G) and VSVdG-Md-CreEGFP(EnvA), approximately 33.5 ml of viral supernatant was layered over 5 ml of 20% (wt/vol) sucrose and concentrated by ultracentrifugation at 21,000 rpm for 2 h at 4 °C in an SW32Ti rotor (Beckman Coulter). Pellets were collected and resuspended in 100 μl DMEM per tube.

Replication-competent VSV was amplified from rescued virus stocks without the use of an amplification plasmid. The medium of confluent BHK-21 cells in a 10-cm dish was replaced with 2 ml DMEM, and 10 μl of supernatant from the rescue well was added to each dish. After 1 h, 13 ml DMEM supplemented with 2% FBS and 1% penicillin-streptomycin was added. Viral supernatant was harvested 2 days later and concentrated by ultracentrifugation at 21,000 rpm for 3 h at 4 °C in an SW32Ti rotor (Beckman Coulter). Pellets were collected and resuspended in 100 μl DMEM per tube.

AAVs were produced as previously described (Xiong et al 2019). Briefly, 293T cells were seeded 20–24 h before transfection onto five poly-D-lysine-coated 15 cm dishes per viral preparation and grown to near confluence. For each preparation, 35 μg AAV Dplasmid (Rep/Cap), 35 μg AAV vector plasmid, and 100 μg pHGTI-adeno1 were mixed in DMEM, combined with 340 μl PEI, and brought to a final volume of 25 ml. At transfection, the medium was replaced with 20 ml DMEM supplemented with 10% NuSerum and 1% penicillin-streptomycin, and 5 ml transfection mixture was added to each dish. Twenty-four hours later, the medium was replaced with 25 ml DMEM.

Three days after transfection, the supernatant was collected and virus was precipitated by addition of PEG-8000 and NaCl to final concentrations of 8.5% (wt/vol) and 0.4 M, respectively, for 2 h at 4 °C. The precipitated virus was pelleted by centrifugation at 7,000 × g for 10 min. The supernatant was discarded, and the pellet was resuspended in lysis buffer (150 mM NaCl, 20 mM Tris, pH 8.0). MgCl2 was added to a final concentration of 1 mM, followed by Benzonase (25 U/ml), and the suspension was incubated at 37 °C for 10–15 min. The virus suspension was then purified by iodixanol gradient ultracentrifugation. Gradients were prepared in OptiSeal tubes by layering 6 ml each of 17%, 25%, 40%, and 60% iodixanol solutions. The virus suspension was loaded onto the gradient, and tubes were centrifuged at 46,500 rpm for 90 min at 16 °C in a Beckman VTi50 rotor. The viral fraction was collected from the 40% iodixanol layer. To remove iodixanol and concentrate the virus, the collected fraction was diluted in PBS and transferred to an Amicon Ultra 100 kDa centrifugal filter unit. The sample was washed three times with PBS by repeated centrifugation at 3,500 rpm for 15 min at 4 °C. The final purified virus was recovered in approximately 150–250 μl PBS.

### Viral vectors

VSVdG-Md-CreEGFP was either coated with native VSV-G or pseudotyped with EnvA (Osakada & Callaway 2013, Wickersham et al 2010). The titer of VSVdG-Md-CreEGFP(VSV-G) was 2.0×10^11^ infectious units per ml (IU/ml). The titer of VSVdG-Md-CreEGFP(EnvA), measured on 293T-TVA800 cells, was 6.9×10^8^ IU/ml and 2.6×10^5^ IU/ml on 293T cells. The titer of replication-competent VSV-Mq-H2BEGFP was 2.2×10^9^ focus-forming units per ml (ffu/ml).

The AAVs used in this study were AAV2/8-Syn-cDIO-VSV-G (6.6×10^14^ genome copies per ml (gc/ml), homemade), AAV2/5-CAG-RABV-N2cP (4.0×10^14^ gc/ml, homemade; N2cP from Addgene plasmid #100808), AAV2/5-CAG-RABV-B19P (4.0×10^14^ gc/ml, homemade; B19P from Addgene plasmid #32631), AAV2/8-Syn-FLPo (1.1×10^13^ gc/ml, purchased from SignaGen Laboratories, SL101451), AAV2/8-fDIO-TVAmCherry (3.6×10^13^ gc/ml, purchased from Salk Viral Vector core), AAV2/8-CAG-TVAmCherry (1.2×10^14^ gc/ml, homemade), and AAV2/9-Syn-cDIO-VSV-G(W72A) (2.3×10^14^ gc/ml, homemade). AAV titers were quantified by quantitative PCR (qPCR) using primers targeting the AAV2 ITR sequence: forward ITR primer, 5’-GGAACCCCTAGTGATGGAGTT-3’ and the reverse ITR primer, 5’-CGGCCTCAGTGAGCGA-3’, following a previously described protocol (Aurnhammer et al 2012).

### Preparation and Stereotaxic injection of AAVs and VSVs

All stereotaxic injections were performed under BSL-2 conditions in the Harvard Center for Comparative Medicine facilities. Mice were anesthetized with a ketamine/xylazine/acepromazine cocktail and placed in a stereotaxic frame. For striatal injections, AAVs and VSVs were injected into either the unilateral dorsal striatum (Figure 1) or the bilateral dorsal striatum (all other figures) at the following coordinates relative to bregma: AP, +1.0 mm; ML, ±1.8 mm; DV, −3.0 mm. AAV helper-virus mixtures were prepared immediately before stereotaxic injection by combining the indicated volumes of each viral stock.

For tracing experiments using VSVdG-Md-CreEGFP(VSV-G), 10 µL of AAV2/8-Syn-cDIO-VSV-G was mixed with 10 µL of AAV2/5-CAG-RABV-N2cP. A total of 250 nL of this mixture was injected into the dorsal striatum. Two weeks later, 100 nL of VSVdG-Md-CreEGFP(VSV-G) was injected into the same region. For tracing experiments using replication-competent VSV-Mq-H2BEGFP, 50 nL of virus was injected into the dorsal striatum.

For tracing experiments using VSVdG-Md-CreEGFP(EnvA), the AAV mixture was prepared by combining 5 µL of AAV2/8-Syn-cDIO-VSV-G, 5 µL of AAV2/5-CAG-RABV-N2cP, 2.5 µL of AAV2/8-Syn-FLPo, and 2.5 µL of AAV2/8-CAG-fDIO-TVA-mCherry. A total of 100 nL of this mixture was injected into the dorsal striatum. Two weeks later, 200 nL of VSVdG-Md-CreEGFP(EnvA) was injected into the same region. For comparison experiments, AAV-RABV-N2cP was either replaced with an equal volume of PBS or AAV-RABV-B19P.

For cytokine-blocking antibody experiments, the nine-cytokine blocking antibody cocktail was prepared by mixing equal volumes of Bio X Cell InVivoMAb™ antibodies targeting IL-6 (BE0046), TNF-α (BE0058), IL-12 (BE0052), IL-18 (BE0237), IL-1α (BE0243), IL-1β (BE0246), CCL2 (BE0185), IL-17A (BE0173), and IL-17F (BE0303). Then, 5 µL of AAV2/8-Syn-cDIO-VSV-G, 5 µL of AAV2/8-CAG-TVA-mCherry, and 10 µL of the antibody cocktail were mixed, and 200 nL of this mixture was injected into the dorsal striatum. Two weeks later, 10 µL of VSVdG-Md-CreEGFP(EnvA) was mixed with 10 µL of the same antibody cocktail, and 400 nL of this mixture was injected into the same region.

For experiments testing fusion-defective VSV-G, 10 µL of AAV2/9-Syn-cDIO-VSV-G(W72A), 5 µL of AAV2/8-CAG-TVA-mCherry, and 5 µL of AAV2/5-CAG-RABV-N2cP were mixed, and 100 nL of this mixture was injected into the dorsal striatum. Two weeks later, 200 nL of VSVdG-Md-CreEGFP(EnvA) was injected into the same region.

For experiments determining the cell types of EGFP-positive cells in the substantia nigra, 10 µl of AAV2/8-Syn-cDIO-VSV-G, 5 µl of AAV2/8-CAG-TVA-mCherry, and 5 µl of AAV2/5-CAG-RABV-N2cP were mixed, and 100 nl of this mixture was injected into the dorsal striatum of IFNAR1/IFNGR1 DKO mice. Two weeks later, 200 nL of VSVdG-Md-CreEGFP(EnvA) was injected into the same region.

### Microglia depletion

To deplete microglia, mice were treated with the CSF1R inhibitor pexidartinib (PLX-3397) throughout the experimental period. In Figure 1, PLX-3397 was administered by oral gavage at 50 mg per kg of mouse body weight. In all other experiments, PLX-3397 was administered through AIN-76A rodent diet (Research Diets Inc., D10001) at 290 mg per kg of diet (Liddelow et al 2017).

### Tissue collection and Immunohistochemistry

Mice were deeply anesthetized and transcardially perfused with phosphate-buffered saline followed by 4% paraformaldehyde (PFA). Brains were post-fixed overnight in 4% PFA and then sequential immersion in 15% and 30% sucrose in PBS at 4 °C until the tissue sank to the bottom of the tube before sectioning. Coronal sections were cut at 25 μm and sagittal sections at 35 μm on a cryostat and collected in serial order onto Superfrost Plus microscope slides.

Immunostaining was performed using a similar general procedure and antibody panel as in our previous study (Krause & Cepko 2024). Sections were washed in PBS for 5 min and blocked for 1 h at room temperature on a shaker in blocking solution consisting of 0.05 M TBS, 1% Triton X-100, and 6% normal donkey serum. After blocking, sections were rinsed twice in 0.05 M TBS and incubated overnight at 4 °C in primary antibody solution containing 0.05 M TBS, 0.2% Triton X-100, and 2% normal donkey serum. Primary antibodies used were rabbit anti-NeuN (1:1000; ab104225, Abcam), rabbit anti-Sox9 (1:300; AB5535, Millipore), rabbit anti-Sox10 (1:300; ab180862, Abcam), and rabbit anti-Iba1 (1:300; GTX100042, GeneTex). After primary incubation, sections were washed twice in 0.05 M TBS for 15 min each and then incubated for 2 h at room temperature on a shaker in secondary antibody solution containing 0.05 M TBS, 0.2% Triton X-100, 2% normal donkey serum, and donkey anti-rabbit Alexa 647 (1:300; 711-605-152, Jackson ImmunoResearch). Sections were then washed three times for 5 min each in 0.05 M TBS and mounted with Fluoromount-G under a coverslip.

### Microscopy and image analysis

For Figure 1, fluorescence images were acquired using a Keyence BZ-X800 fluorescence microscope equipped with a 10× objective. Images were merged using BZ-X800 Analyzer software. All other images were acquired on Nikon microscopes using NIS-Elements AR software.

For representative images, widefield fluorescence images were acquired as stitched fields using either a Plan Apo λ 20× objective (NA 0.75) or a Plan Apo 10× DIC objective (NA 0.45).

To quantify viral spread in the substantia nigra, coronal sections were cut at 25 μm, collected serially, and analyzed at approximately 200-μm intervals. Z-stack images were acquired from 4–5 coronal sections per brain using a Plan Apo λ 20× objective (NA 0.75). Z-stacks were collected as 24 optical sections at 0.9-μm intervals. Images were processed in ImageJ/Fiji using maximum-intensity z projection, and EGFP-positive cells were counted manually. Values from the left and right sides were averaged to generate a single data point for each mouse.

To identify starter cells, z-stack images were acquired using a Plan Apo λ 20× objective (NA 0.75), with 13 optical sections collected at 0.9-µm intervals. Starter cells were defined as EGFP+ / mCherry+ double-positive cells. Colocalization was assessed manually by examining signal overlap within single optical section and in orthogonal views of the z-stack.

To determine the cell types of VSV-spread cells in substantia nigra, sections were immunostained and imaged using a Plan Apo λ 20× objective (NA 0.75), and z-stacks were acquired as 18 optical sections at 0.6-μm intervals. Colocalization of EGFP-positive cells with far-red immunoreactive signals for NeuN, Sox9, Sox10, or Iba1 was assessed manually based on signal overlap within single optical sections and in orthogonal views of the z-stack.

### Data and Resource Availability

Representative high-resolution microscopy images and associated metadata will be deposited in a public data repository upon publication. Source data underlying the quantitative analyses are provided in the Supplementary Materials. All plasmids generated in this study will be deposited to Addgene. Where feasible, we will also pursue future distribution of viral reagents through a shared research core, such as Salk Institute’s Viral Vector Core (GT3).

### Statistical analysis

Data are presented as mean ± SEM. Statistical analyses were performed using GraphPad Prism. Sample sizes are shown in the figures. Normality was assessed using the Shapiro–Wilk test, and homogeneity of variances was evaluated using the Brown– Forsythe test. Because both assumptions were satisfied, one-way ANOVA followed by Dunnett’s multiple comparisons test was used in the analysis. Significance thresholds: ns (not significant) *p* ≥ 0.05, *p* < 0.05.

## Results

### Requirements for One-Step Anterograde Tracing with VSVdG coated by VSV-G

In the RABV retrograde tracing systems, the RABV-G is typically expressed in the starter cells prior to infection with the tracing virus (Callaway & Luo 2015, Wickersham et al 2007b). Due to the toxicity of VSV-G, we could not use this strategy, and thus delayed expression of VSV-G, turning it on by infection with the tracing virus. This was accomplished by generating a VSVdG variant encoding CreEGFP and using a Cre-dependent AAV to express VSV-G, only after VSVdG-CreEGFP infected the starter cells. In this approach, Cre/Flp-intersectional AAVs would be required for cell-type-specific VSV-G expression (Fenno et al 2014, Hafner et al 2019). A second adaptation to tracing with VSVdG was to reduce toxicity due to the VSV-M protein. M induces toxicity by shutting off host gene expression. VSV-M mutants have been identified that reduce this host shut-off (Ahmed et al 2003, Hoffmann et al 2010, Jayakar & Whitt 2002, Stojdl et al 2003). A VSVdG-CreEGFP variant was thus created with such mutations, here the M33A/M51R mutations, “VSVdG-Md-CreEGFP”. One further adaption was made due to the fact that attenuating VSV-M–mediated host shut-off was expected to allow the host to produce antiviral responses. These responses would reduce the ability of VSVdG-Md-CreEGFP to spread. As interferon has been shown to inhibit VSV replication, and AAV can induce other cytokines that might as well, we tested multiple strategies to disrupt cytokine production and downstream signaling pathways. We ultimately found that, in mice lacking both type I and type II interferon receptors, a combination of AAV-mediated expression of RABV-P protein and microglia depletion was sufficient to enable efficient one-step anterograde tracing by AAV-mediated trans-complementation of VSVdG (Figure 1B).

The AAVs encoding VSV-G and RABV-P and VSVdG were injected into the dorsal striatum of IFNAR1/IFNGR1 double-knockout (DKO) mice. The AAVs used were AAV2/8-Syn-cDIO-VSV-G and AAV2/5-CAG-RABV-P. RABV-P was derived from the CVS-N2c strain (Reardon et al 2016). AAV2/5 was used for delivery of RABV-P because this AAV serotype has been reported to produce broader and more efficient transduction in the dorsal striatum than several other commonly used serotypes (Aschauer et al 2013). In addition to the delivery of these AAVs and the use of the IFNAR1/IFNGR1 DKO mice, animals also had microglia depletion. The colony-stimulating factor 1 receptor (CSF1R) kinase inhibitor, pexidartinib (PLX3397), was used to deplete microglia throughout the entire experimental period (Elmore et al 2014, Li et al 2017, Liddelow et al 2017) (Figure 1A-B). Two weeks after injection with the AAVs, VSVdG was delivered to the dorsal striatum, and five to seven days later, viral spread was assessed in the substantia nigra, a region distant from the injection site (Figure 1A). The dorsal striatum is organized as a mosaic of two distinct compartments, patch and matrix. Projection neurons in the striatal patch project preferentially to the dorsal, cell-dense substantia nigra pars compacta (SNc), whereas striatal matrix neurons project to the ventral, cell-sparse substantia nigra pars reticulata (SNr) (Fujiyama et al 2011, Gerfen 1984, Gerfen 1992, McGregor et al 2019, Watabe-Uchida et al 2012). The striatal infection by VSVdG would not discriminate between patch and matrix neurons, and thus VSV spread would be expected to both downstream target regions. Indeed, this was observed as EGFP-labelled cells were observed in the SNc and SNr. Moreover, we observed VSVdG-Md labeling in the ventral tegmental area (VTA) (Figure 1C). Although the dorsal striatum–VTA pathway is less well characterized, previous monosynaptic retrograde rabies-tracing studies have identified direct inputs from the dorsal striatum to the VTA (Beier et al 2019, Beier et al 2015, Faget et al 2016, Watabe-Uchida et al 2012).

The experiments described above provided evidence that VSVdG spread could be achieved using a combination of methods that block innate immune responses. In order to further dissect the necessity and efficacy of different methods to block antiviral signaling pathways, additional experiments were done. In IFNAR1/IFNGR1 DKO mice, we either omitted microglia depletion or removed the AAV-RABV-P. In addition, immunocompetent C57BL/6 mice and several immunodeficient mouse lines were tested with microglia depletion and AAV-RABV-P. Because both AAV and VSV can be sensed by TLRs and RLRs, we tested mice lacking the major downstream adaptor proteins for these pathways. Specifically, MyD88 knockout and TRIF loss-of-function mutant mice were examined because they mediate TLR signaling, whereas a MAVS knockout mice was examined because it mediates RLR signaling (Chan et al 2021, Georgel et al 2007, Hosel et al 2012, Jiang et al 2005, Kato et al 2005, Kawai & Akira 2009, Lund et al 2004, Shao et al 2018, Shi et al 2011, Yoneyama et al 2004, Zhu et al 2009). Single knockouts of Myd88, TRIF, or MAVS have been shown to confer strong susceptibility to VSV in some cases, whereas in other cases, triple knockouts were required (Gern et al 2024, Ghita et al 2021, Lang et al 2007, Pavlou et al 2025, Spanier et al 2014, Sun et al 2006, Zhou et al 2007) (Figure 1D). Therefore, we compared mouse lines missing these adaptors with IFNAR1/IFNGR1 DKO mice. Together, these experiments revealed that, in IFNAR1/IFNGR1 DKO mice, both microglia depletion and AAV-RABV-P were required for successful viral spread. In contrast, neither immunocompetent C57BL/6 mice nor any of the other immunodeficient strains showed efficient spread, in the presence of both microglia depletion and AAV-RABV-P (Figure 1C). Interestingly, RABV-P has been shown to inhibit type I interferon induction and downstream signaling pathways shared by both type I and type II interferons (Brzozka et al 2005, Brzozka et al 2006, Vidy et al 2005, Vidy et al 2007). Its requirement in IFNAR1/IFNGR1 DKO mice suggests that RABV-P suppresses additional antiviral pathways that remain active in these mice, possibly including responses mediated by other pro-inflammatory cytokines.

The experiments described above examined VSVdG spread to areas some distance from the dorsal striatum. We also evaluated local spread within the dorsal striatum and examined the effects of blocking innate immunity locally. This local spread likely occurred through collateral axons of striatal neurons (Burke et al 2017). In IFNAR1/IFNGR1 DKO mice, we compared four conditions: VSVdG-Md alone as a control, VSVdG-Md with AAV-VSV-G together with either AAV-RABV-P, microglia depletion, or both. Intrastriatal spread was observed only in groups receiving both AAV-VSV-G and AAV-RABV-P, with or without microglia depletion. Mice receiving AAV-VSV-G with microglia depletion alone, i.e. without AAV-RABV-P, did not show intrastriatal spread (Figure 1C). Thus, RABV-P is sufficient to overcome barriers to local VSVdG-Md spread in IFNAR1/IFNGR1 DKO mice, whereas long-distance spread requires additional microglia depletion.

As the protocol for VSVdG spread employs transduction with AAV to supply VSV-G, it was of interest to determine whether the restrictive immune responses were triggered by VSV infection alone, or along with AAV transduction. Because replication-competent VSV encodes its own VSV-G, it does not require AAV-mediated VSV-G complementation and therefore allowed us to assess how VSV infection alone is restricted across wild-type and immunodeficient mouse strains. Previous *in vivo* studies in wild-type mice showed that intracerebral injection of replication-competent VSV carrying the host-shutoff-defective M51 deletion in VSV-M remains lethal (Jiang et al 2021, Lun et al 2006, Muik et al 2014), whereas VSV carrying the more attenuated VSV-Md variant still causes mild paralysis (Clarke et al 2007, Cooper et al 2008). Because the VSV-Mq variant is more attenuated in its host-shutoff activity than the Md variant (Hoffmann et al 2010), we generated a mutant virus carrying the VSV-Mq variant to determine whether replication-competent VSV spread could be evaluated across immunodeficient mouse strains despite the potential for severe viral pathogenicity. The virus also encoded H2BEGFP to label the nuclei of infected cells.

We injected replication-competent VSV-Mq-H2BEGFP into the dorsal striatum of C57BL/6, IFNAR1-knockout, IFNGR1-knockout, and MAVS-knockout mice. VSV spread was assessed 3 days later because the health status of IFNAR1-knockout mice deteriorated rapidly (Figure S1A). In C57BL/6 mice, VSV-infected cells were detected only around the injection site. This restricted pattern contrasts with our previous findings using replication-competent VSV carrying wild-type VSV-M, in which striatal injection led to viral spread to several additional brain regions, including the substantia nigra, at the same post-injection time point (Drokhlyansky et al 2017, Mundell et al 2015). Similarly, in IFNGR1-knockout mice, VSV-infected cells were detected only around the injection site. In contrast, in IFNAR1-knockout mice, VSV spread across a much broader region of the forebrain, including the cortex. In MAVS-knockout mice, VSV also spread to multiple forebrain regions, although to a lesser extent. In both IFNAR1- and MAVS-knockout mice, the distribution of infected cells was consistent with multistep spread following VSV infection. However, neither knockout was sufficient to permit long-distance spread to the substantia nigra (Figure S1B). These results suggest that the MAVS–IFNAR axis is the major pathway by which VSV infection is sensed and restricted in the brain (Figure 1D).

Importantly, IFNAR1 knockout alone was sufficient to support multistep spread of VSV-Mq-H2BEGFP in the absence of AAV transduction. In contrast, in the one-step anterograde tracing system, AAV-RABV-P was still required to support local spread of VSVdG-Md in IFNAR1/IFNGR1 DKO mice (Figure 1C). This difference suggests that prior AAV transduction may activate additional antiviral pathways that are independent of type I and type II interferon signaling and continue to restrict VSVdG-Md spread. The ability of RABV-P to restore local VSVdG-Md spread indicates that RABV-P can overcome this remaining antiviral inhibition. However, the failure of VSV-Mq-H2BEGFP to spread to the substantia nigra in either IFNAR1- or MAVS-knockout mice further indicates that disruption of the MAVS–type I interferon pathway is not sufficient to permit long-distance VSV propagation. These barriers may include non-interferon pro-inflammatory cytokines released by microglia. Consistent with this possibility, wild-type VSV-M broadly suppresses host gene expression, including interferons and pro-inflammatory cytokines, which may allow replication-competent VSV to spread both locally and to distant brain regions (Drokhlyansky et al 2017, Mundell et al 2015). In contrast, host-shutoff-defective VSV-M variants, including VSV-Md in VSVdG-Md and VSV-Mq in replication-competent VSV-Mq-H2BEGFP, may permit continued inflammatory gene expression that restricts long-distance spread. This model may explain why IFNAR1/IFNGR1 DKO combined with RABV-P expression is sufficient to support local VSVdG-Md spread but not spread to the substantia nigra. The requirement for additional microglia depletion suggests that microglia-mediated inflammatory responses represent a major barrier to long-range VSVdG-Md propagation.

### Requirements for One-Step Anterograde Tracing with VSVdG pseudotyped by EnvA

In order to further develop VSV for monosynaptic anterograde tracing, it was necessary to target infection to specified starter cells. This was achieved by pseudotyping the tracing virus with EnvA, the envelope glycoprotein from avian sarcoma and leukosis virus, and selectively expressing its receptor, TVA, in the intended starter cells (Callaway & Luo 2015, Wickersham et al 2007b). We generated EnvA-pseudotyped VSVdG-Md-CreEGFP (Figure S2) and evaluated the adaptations described above for one-step anterograde tracing from defined starter cells. In addition to AAV2/8-Syn-cDIO-VSV-G and AAV2/5-CAG-RABV-P, two additional AAVs were injected, AAV2/8-Syn-FLPo and AAV2/8-CAG-fDIO-TVAmCherry, into the dorsal striatum (Figure 2B). In this protocol, striatal neurons, including both Drd1+ and Drd2+ medium spiny neurons (MSNs), were designated starter cells and expressed TVA. Two weeks later, VSVdG-Md-CreEGFP(EnvA) was injected into the same region, and VSV spread was assessed 6 days later. We examined all three major direct projection targets of the dorsal striatum: the globus pallidus pars externa (GPe), globus pallidus pars interna (GPi), and substantia nigra (Nelson & Kreitzer 2014) (Figure 2A). Moreover, we quantified EGFP+ cell numbers in the substantia nigra because its well-defined boundaries and distance from the injection site enabled reliable analysis. As expected, in IFNAR1/IFNGR1 DKO mice treated with PLX-3397 throughout the experimental period, VSV spread to all three regions (Figure 2C).

**Figure 2.**
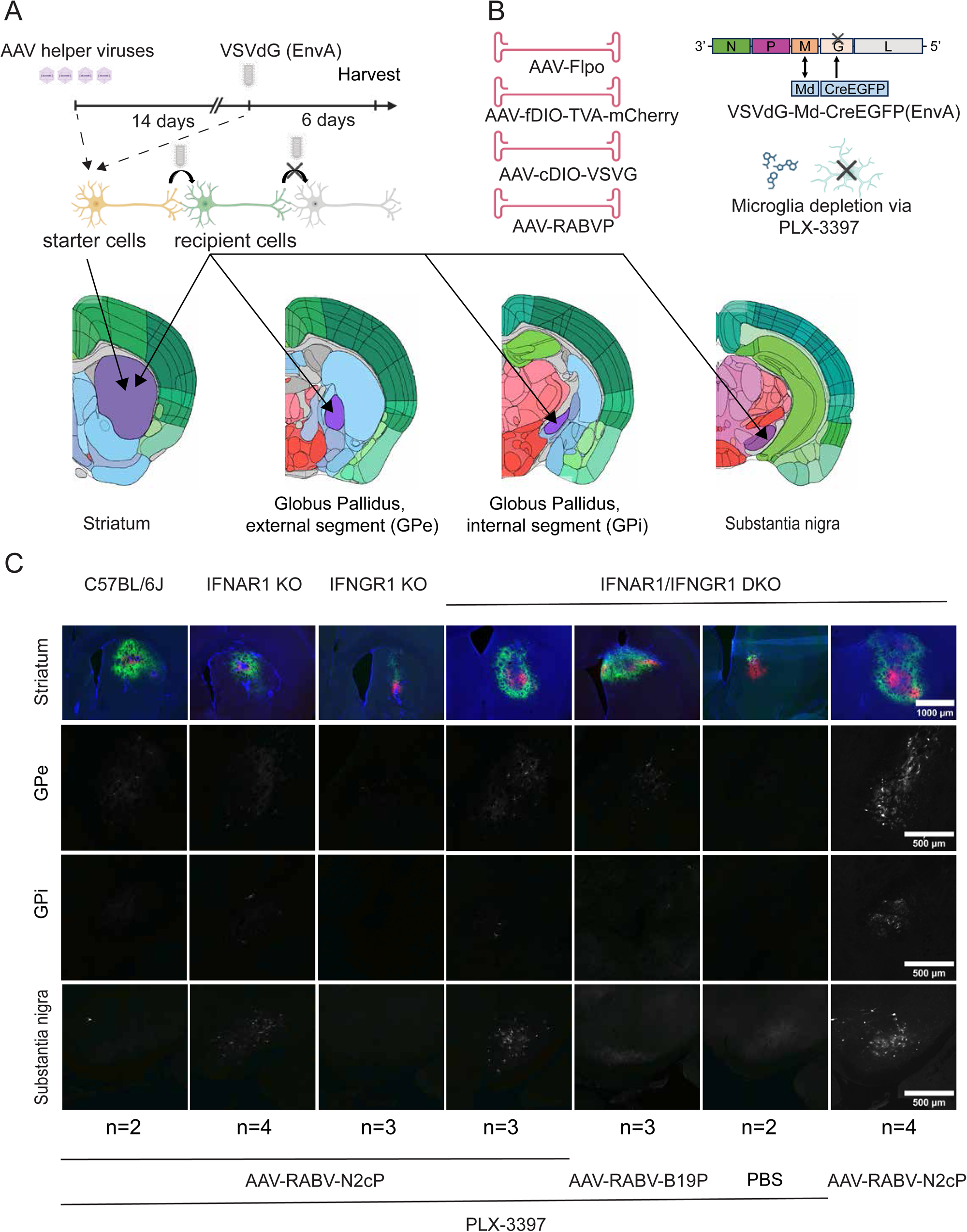
Requirements for one-step anterograde tracing with VSVdG-Md-CreEGFP(EnvA) (A) Experimental design. (B) The AAVs, VSV, and microglia-depletion drug used in this experiment. (C) Representative images of coronal sections showing the dorsal striatum and three downstream target regions under the indicated conditions. In the dorsal striatum, both mCherry labeling after AAV-TVA-mCherry transduction and EGFP labeling after VSV infection are shown. In the three downstream target regions, only EGFP labeling after VSV infection is shown. In IFNAR1/IFNGR1 DKO mice, four conditions were tested: AAV-RABV-N2cP alone, microglia depletion alone, AAV-RABV-N2cP plus microglia depletion, and AAV-RABV-B19P plus microglia depletion. In contrast, in C57BL/6, IFNAR1-knockout, and IFNGR1-knockout mice, only the AAV-RABV-N2cP plus microglia depletion condition were tested. The number of mice in each group is indicated below the corresponding image.

We next tested whether microglia depletion, AAV-RABV-P, and IFNAR1/IFNGR1 DKO were each required in this context. Unexpectedly, VSV spread to all three downstream brain regions even in the absence of microglia depletion. Quantification of EGFP+ cells in the substantia nigra showed that VSV spread was unchanged by microglial depletion (Figure 3D, E). This result contrasted with the condition in which VSVdG was coated with VSV-G, where microglia depletion was required for long-distance spread. Because VSV-G present on the VSV virion activates TLR4/CD14-dependent innate immune signaling (Georgel et al 2007), and microglia are the principal CNS cell type with high constitutive expression of both TLR4 and CD14 (Zhang et al 2014), we hypothesized that EnvA-pseudotyped VSVdG does not induce microglial antiviral cytokine release to the same extent as VSV-G-coated VSVdG, thereby permitting long-distance spread even in the presence of microglia.

**Figure 3.**
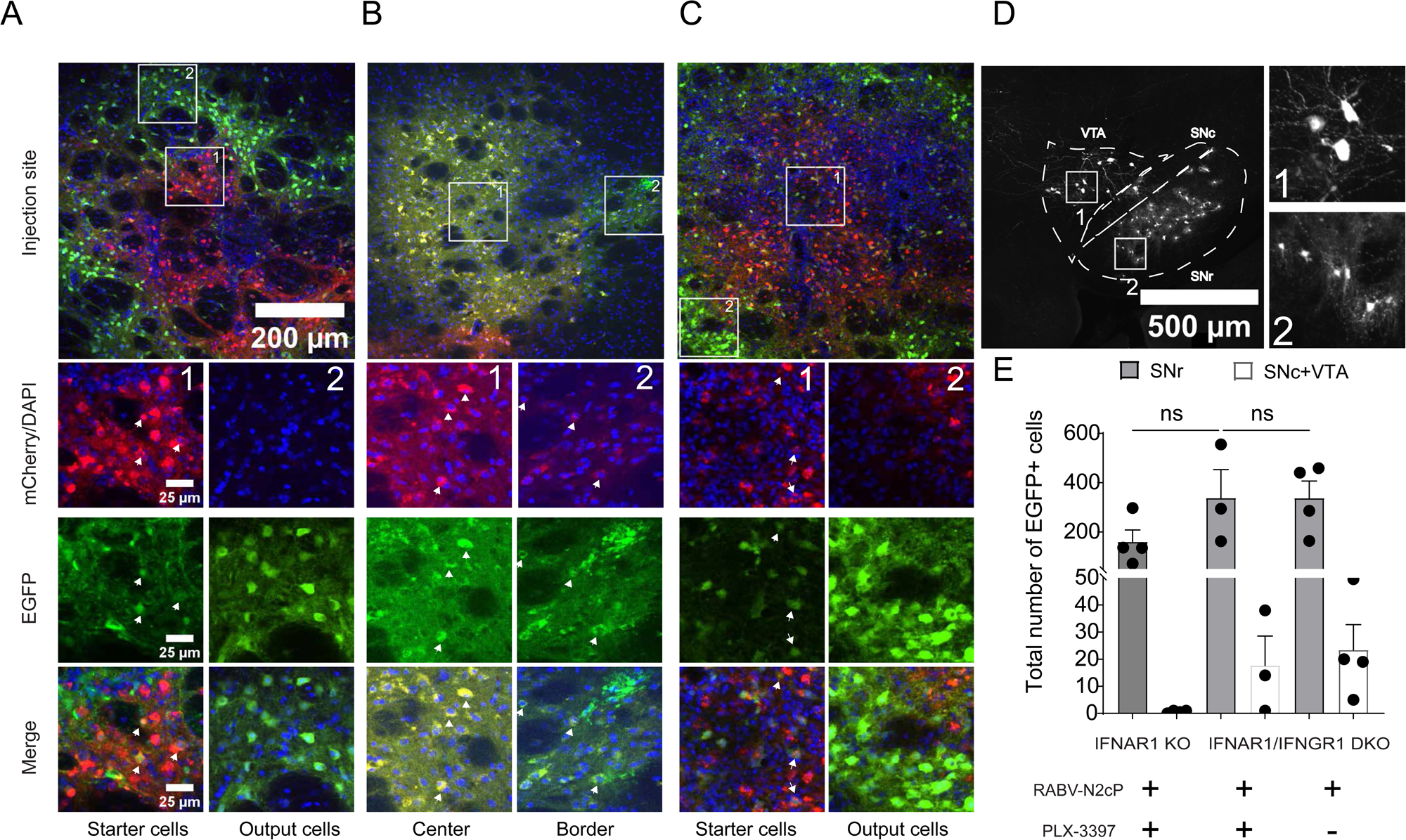
Starter cells in the dorsal striatum and long-distance VSV spread during one-step anterograde tracing with VSVdG-Md-CreEGFP(EnvA) (A) Representative image of putative starter cells in the dorsal striatum, identified by co-labeling for mCherry and EGFP, from IFNAR1/IFNGR1 DKO mice under the AAV-RABV-N2cP plus microglia depletion condition. EGFP-only cells located at the striatal border are shown for comparison. n = 3. (B) Representative images of putative starter cells in the dorsal striatum, from IFNAR1/IFNGR1 DKO mice under the AAV-RABV-N2cP plus microglia depletion condition. Animals received all AAVs except AAV2/8-Syn-cDIO-VSV-G. n=3. (C) Representative image of putative starter cells in the dorsal striatum, identified by co-labeling for mCherry and EGFP, from IFNAR1/IFNGR1 DKO mice under the AAV-RABV-N2cP alone condition. EGFP-only cells located at the striatal border are shown for comparison. n = 4. (D) Representative images showing EGFP labeling after VSV spread to the substantia nigra and VTA in IFNAR1/IFNGR1 DKO mice under the AAV-RABV-N2cP alone condition. n=4. (E) Quantification of EGFP-positive cells in the SNr and in the combined SNc/VTA region.

To assess the requirement for AAV-mediated expression of RABV-P from the CVS-N2c strain (AAV-RABV-N2cP), we either omitted this AAV or replaced it with an AAV expressing RABV-P from the vaccine-derived SAD B19 strain (AAV-RABV-B19P), with microglia depletion. RABV of the SAD B19 strain has been widely used for monosynaptic retrograde tracing, however, the CVS-N2c strain has been shown to support more efficient transsynaptic spread (Callaway & Luo 2015, Reardon et al 2016, Wickersham et al 2007b, Xu et al 2020). Consistent with the results from one-step anterograde tracing using VSV-G-coated VSVdG-Md (Figure 1C), AAV-RABV-N2cP was required for VSV spread to the substantia nigra (Figure 2C). When AAV-RABV-P was omitted, VSV spread was not detected in downstream targets of the dorsal striatum. In mice receiving AAV-RABV-B19P, VSV spread was observed mainly in the GPe, with minimal spread to the substantia nigra (Figure 2C). These results reveal strain-specific differences in the ability of RABV-P to overcome the antiviral effects of cytokines that restrict VSV spread in one-step anterograde tracing. They further suggest that, in addition to RABV-G (Kim et al 2016), RABV-P may be another strain-specific factor contributing to the efficiency of monosynaptic retrograde tracing.

To assess the requirement for IFNAR1/IFNGR1 DKO, we compared IFNAR1/IFNGR1 DKO mice with C57BL/6, IFNAR1-knockout, and IFNGR1-knockout mice in the presence of microglia depletion and AAV-RABV-N2cP. We found that VSVdG spread was minimal in C57BL/6 and IFNGR1 knockout mice, whereas in IFNAR1 knockout mice VSVdG spread efficiently to all three downstream targets of the dorsal striatum. Quantification of EGFP-positive cells in the SNr showed a trend toward reduced labeling in IFNAR1 knockout mice compared with IFNAR1/IFNGR1 DKO mice, although the difference was not statistically significant (Figure 3D, E, Data S1). We also quantified EGFP-positive cells in regions adjacent to the SNr, including the SNc and VTA. Labeling in these regions was minimal in IFNAR1 knockout mice but readily detected in IFNAR1/IFNGR1 DKO mice, suggesting that IFN gamma signaling also restricts the efficiency of VSVdG spread. Moreover, consistent with the established organization of striatal outputs (Gerfen 1985), labeling was substantially greater in the SNr than in the SNc.

In the dorsal striatum, putative starter cells were identified by co-labeling for mCherry and EGFP, derived from AAV-TVA-mCherry and VSVdG-Md-CreEGFP(EnvA), respectively. In IFNAR1/IFNGR1 DKO mice receiving AAV-RABV-N2cP, with (Figure 3A) or without (Figure 3C) microglia depletion, mCherry-positive putative starter cells showed weaker EGFP signal and compromised neuronal morphology compared with EGFP-only cells near the striatal border. These EGFP-only cells were presumably infected by viral spread from starter cells. The unhealthy morphology of putative starter cells could have been caused by VSVdG-Md replication, VSV-G expression, or injury from intrastriatal injection. To distinguish among these possibilities, we examined IFNAR1/IFNGR1 DKO mice that received AAV-RABV-N2cP and microglia depletion but not AAV-VSV-G. Under this condition, most EGFP-positive cells in the dorsal striatum were also mCherry-positive. This result is consistent with infection being restricted mainly to starter cells, with little or no viral spread in the absence of VSV-G. A few EGFP-only cells were still observed. These cells may have expressed TVA-mCherry below the detection threshold but at levels sufficient for EnvA-mediated infection, because the EnvA–TVA interaction is highly sensitive and very low levels of TVA can support initial infection (Seidler et al 2008, Wall et al 2010). Importantly, even without VSV-G expression, neurons showed unhealthy morphology in both the central dorsal striatum near the injection site and the striatal border region farther from the injection site (Figure 3B). Thus, neuronal stress was not confined to the injection site and did not require VSV-G expression. These results suggest that VSVdG-Md replication itself in starter cells can compromise neuronal health in this context.

We further investigated RABV-P–dependent immune antagonism by replacing AAV-RABV-N2cP with a cocktail of nine antibodies targeting pro-inflammatory cytokines. We focused on cytokines reported to be induced by AAV transduction or VSV infection or involved in virus-associated inflammatory responses (Chan et al 2021, Hodges et al 2001, Rajan et al 2011, Rogers et al 2011, Sahu et al 2021, Schreiber et al 2019, Smedberg et al 2014, Smith et al 2022, Xiong et al 2019). Because type I and type II interferon signaling were addressed genetically using IFNAR1/IFNGR1 DKO mice, we selected nine commercially available Bio X Cell InVivoMAb™ antibodies targeting IL-6, TNF-α, IL-12, IL-18, IL-1α, IL-1β, CCL2, IL-17A, and IL-17F. Several of these cytokines, including TNF-α and IL-12, have been reported to inhibit productive VSV replication, in some cases synergistically with interferons (Belkowski & Sen 1987, Chauhan et al 2010, Komatsu et al 1999, Maheshwari & Friedman 1979, Mariani et al 2019, Mestan et al 1988, Muller et al 1994, Trottier et al 2005, Tsukamoto & Price 1982, Wong & Goeddel 1986). The antibody mixture was co-injected with both the AAVs and VSVdG-Md-CreEGFP (EnvA) (Figure 4A). To increase the concentrations of both antibody and AAV in the injection mixture, the combination of AAV2/8-Syn-FLPo and AAV2/8-CAG-fDIO-TVA-mCherry was replaced with a single AAV2/8-CAG-TVA-mCherry vector (Figure 4B). We found that in IFNAR1/IFNGR1 DKO mice, this antibody cocktail was sufficient to enable intra-striatal VSVdG-Md spread, as well as spread to all three downstream projection targets of the dorsal striatum (Figure 4C, D). We also noted that the single AAV2/8-CAG-TVA-mCherry vector likely labeled both neurons and glial cells as starter cells (Hollidge et al 2022), whereas the combination of AAV2/8-Syn-FLPo and AAV2/8-CAG-fDIO-TVA-mCherry restricted labeling of starter cells to primarily neurons. This suggests that, even if VSVdG-Md-CreEGFP(EnvA) initially enters glial cells, broad expression of AAV2/5-CAG-RABV-N2cP is sufficient to overcome antiviral responses triggered by these cells.

**Figure 4.**
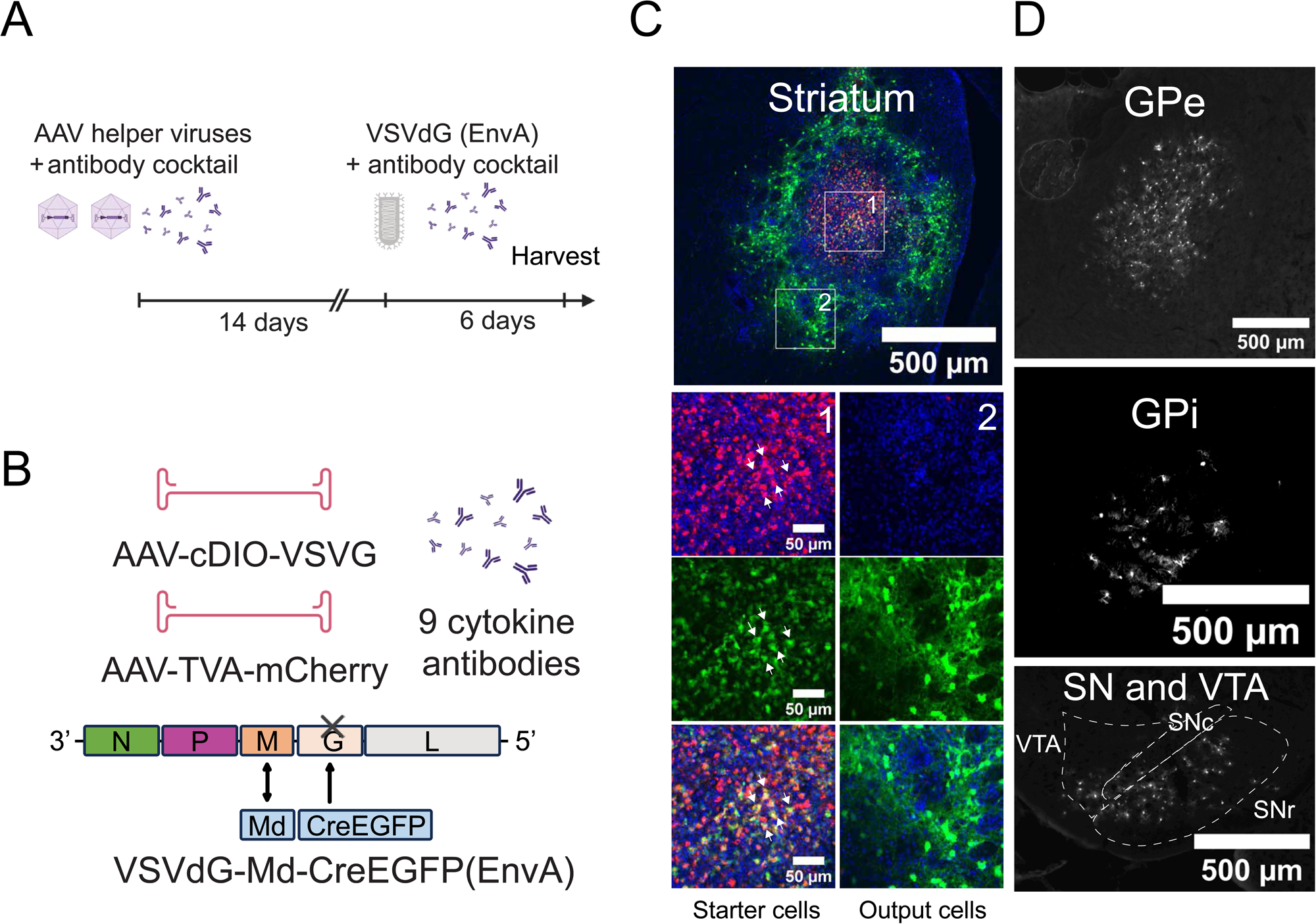
Test of whether a cytokine-blocking antibody cocktail can replace AAV-RABV-N2cP to enable one-step anterograde tracing with VSVdG-Md-CreEGFP(EnvA) (A) Experimental design. (B) AAVs, VSV, and cytokine-blocking antibody cocktail used in this experiment. (C) Representative image of putative starter cells in the dorsal striatum, identified by co-labeling for mCherry and EGFP. EGFP-only cells located at the striatal border are shown for comparison. (D) Representative images of three downstream projection targets of the dorsal striatum. EGFP labeling after VSV infection is shown. n = 2.

After establishing that EnvA-pseudotyped VSVdG spread to the expected brain regions following striatal injection with AAVs, a series of coronal sections and one sagittal section were examined to determine whether viral spread was restricted to predicted projection targets. The coronal sections were obtained from IFNAR1/IFNGR1 DKO mice that received AAV-RABV-N2cP with microglia depletion, whereas the sagittal section was obtained from IFNAR1/IFNGR1 DKO mice that received AAV-RABV-N2cP without microglia depletion. In both conditions, EGFP+ cells were detected only in the expected basal ganglia projection targets, together with mCherry+ axons, and were not observed in any other brain regions (Figure 5A, B). These results indicate that VSVdG spread was confined to the predicted direct striatal output regions, consistent with one-step anterograde spread only.

**Figure 5.**
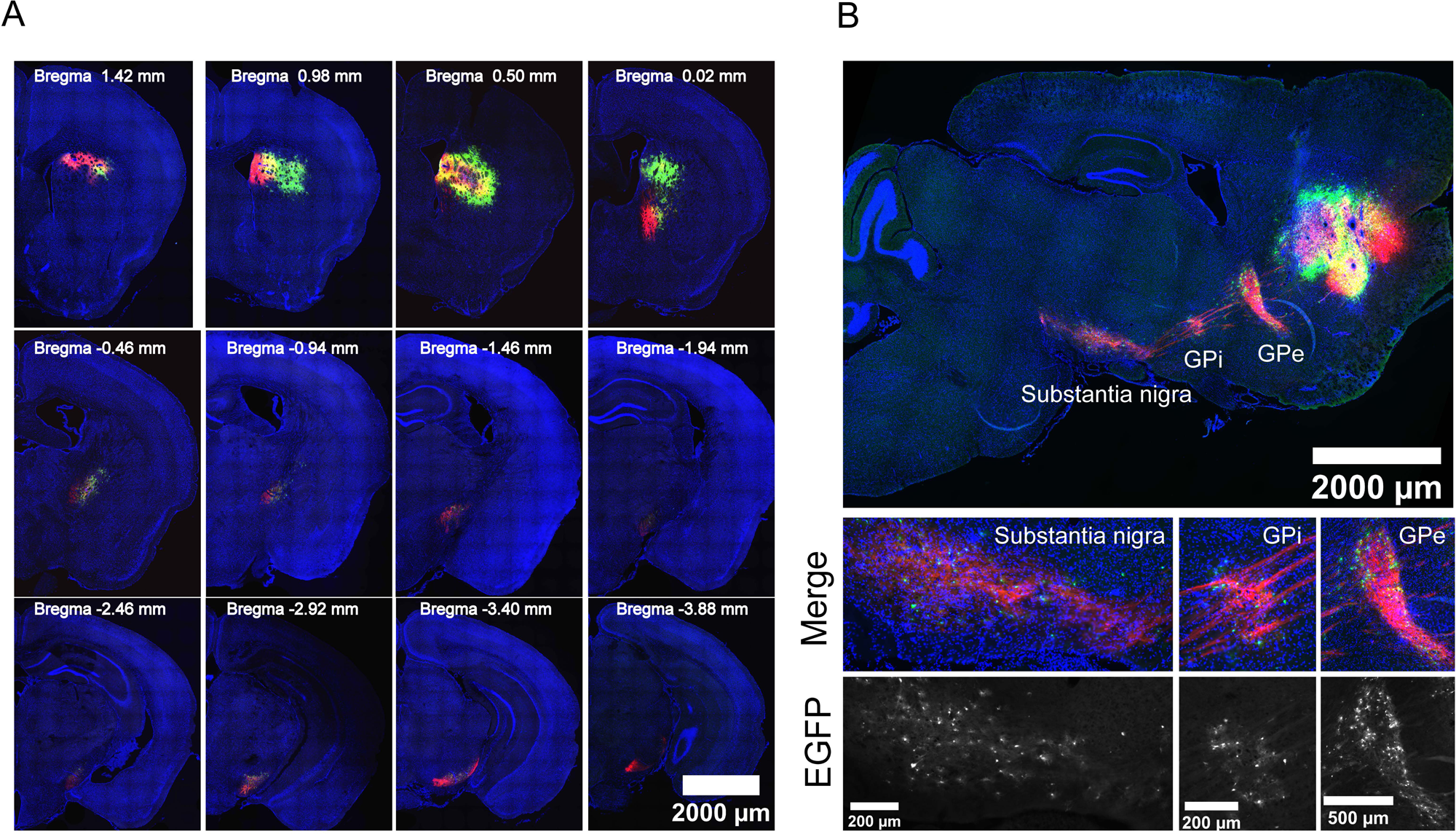
Whole-brain spread pattern of VSV in coronal and sagittal sections (A) Serial coronal sections from IFNAR1/IFNGR1 DKO mice receiving AAV-RABV-N2cP with microglia depletion are shown from anterior to posterior, with approximate Bregma coordinates assigned by comparison with a mouse brain stereotaxic atlas (Paxinos & Franklin 2003). mCherry labeling from AAV-TVA-mCherry transduction and EGFP labeling from VSV infection are shown. (B) Sagittal section from an IFNAR1/IFNGR1 DKO mouse receiving AAV-RABV-N2cP alone. Both mCherry labeling after AAV-TVA-mCherry transduction and EGFP labeling after VSV infection are shown.

### VSVdG Spreads to Both Neurons and Glial Cells in One-Step Anterograde Tracing

Monosynaptic viral tracing is intended to label synaptically connected neurons (Callaway & Luo 2015). It was thus important to determine whether the EGFP+ cells in the downstream projection targets of the dorsal striatum were neurons only. NeuN immunohistochemistry was performed on sections from IFNAR1/IFNGR1 DKO mice transduced with AAV-RABV-N2cP without microglia depletion. Mice were infected with VSVdG-Md-CreEGFP(EnvA) two weeks later and harvested 6 days after VSV injection. NeuN immunostaining was broadly reduced across the GPe and substantia nigra compared with neighboring regions that did not receive striatal projections. This reduction was not confined to EGFP-positive VSVdG-infected cells, and NeuN/EGFP colocalization was rarely observed (Figure 6A). Because these regions receive axonal projections from starter neurons expressing VSV-G, we examined whether the fusogenic activity of VSV-G contributed to the reduced NeuN immunoreactivity. To test this possibility, we used a fusion-defective VSV-G variant carrying the W72A mutation in the fusion loop (Stanifer et al 2011). Three AAVs, AAV2/5-CAG-RABV N2cP, AAV2/8-CAG-TVA-mCherry and AAV2/9-Syn-cDIO-VSV-G(W72A) were injected into the dorsal striatum. Two weeks later, VSVdG-Md-CreEGFP (EnvA) was injected into the same region, and brain samples were collected 6 days later (Figure 6B), matching the time point used in Figure 6A. Under this condition, we did not detect EGFP signal resulting from VSVdG spread in the substantia nigra, consistent with the requirement for VSV-G fusion activity in VSV infection and spread. However, NeuN immunostaining showed clear labeling in substantia nigra, and NeuN+ cells were less densely distributed in the SNr than in the SNc, consistent with previous studies (Figure 6C) (Erb et al 2024, Koprich et al 2010, Zheng et al 2021). These results suggest that reduced NeuN immunoreactivity in striatal projection regions reflects neuronal stress caused by VSV-G fusogenic activity at axon terminals of starter neurons.

**Figure 6.**
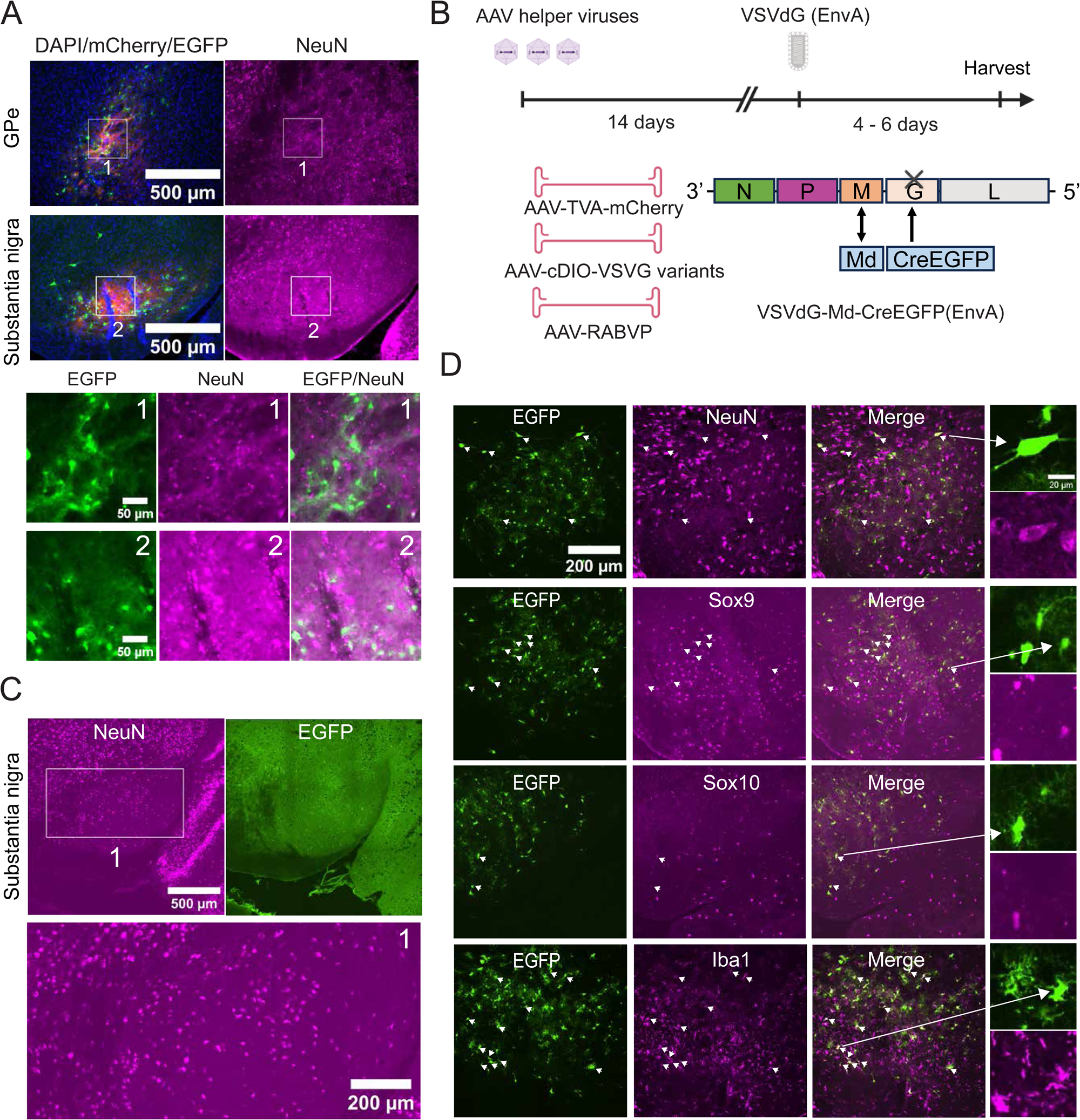
Determination of whether EGFP-positive cells in the substantia nigra are neurons or glial cells (A) Representative images of NeuN immunostaining in downstream projection targets from IFNAR1/IFNGR1 DKO mice receiving AAV-RABV-N2cP alone. NeuN immunostaining and EGFP labeling after VSV infection are shown in the GPe and substantia nigra. (B) Experimental design and AAVs and VSV used in this experiment. (C) Representative images from mice receiving AAV expressing fusion-defective VSV-G W72A. NeuN immunostaining and EGFP-channel images used to assess EGFP-positive cells in the substantia nigra are shown. n=2. (D) Representative images used to assess colocalization of EGFP with NeuN, SOX9, SOX10, and Iba1 in the substantia nigra at 4 days after VSV injection. n=2.

To further interrogate the identity of EGFP+ cells in the target regions, immunohistochemistry for glial cell markers were used. AAV2/9-Syn-cDIO-VSV-G(W72A) was replaced with AAV2/8-Syn-cDIO-VSV-G wild type, while keeping the other two AAVs unchanged. These AAVs were injected into the dorsal striatum, followed two weeks later by injection of VSVdG-Md-CreEGFP (EnvA) into the same region. Brain samples were collected 4, 5, or 6 days after VSV injection (Figure 6B). NeuN labeling was best preserved at 4 days post VSV infection, so this time point was chosen for further analysis. Immunohistochemistry with the astrocyte marker, SOX9, the oligodendrocyte-lineage marker, SOX10, and the microglia marker, Iba1, was carried out. EGFP+ cells colocalized with each of these four markers, suggesting that VSVdG spread to both neurons and glial cells in the substantia nigra (Figure 6D).

## Discussion

Although replication-competent VSV has previously been used for multistep anterograde tracing, simply combining VSVdG with an AAV expressing VSV-G was insufficient to achieve one-step anterograde spread. We reasoned that prolonged VSV-G expression and VSV M protein–mediated host shutoff could compromise cell viability. We therefore generated VSVdG-Md-CreEGFP, which carries the attenuating M33A/M51R mutations in the M protein (VSV-Md), and combined it with a Cre-dependent AAV-VSV-G vector to restrict VSV-G expression to a short time window following VSV infection. Although these mutations reduced M-mediated cytotoxicity and host shutoff, they also weakened the virus’s ability to suppress innate antiviral responses, thereby creating an additional barrier to VSVdG-Md spread. Here, we investigated the host antiviral mechanisms restricting VSVdG-Md propagation using AAV-mediated expression of rabies virus phosphoprotein, mouse models deficient in innate immune signaling pathways, pharmacological depletion of microglia, and cytokine-blocking antibody cocktail.

Our findings demonstrate that type I interferon signaling is a dominant barrier to VSVdG-Md spread. Type I interferon may inhibit VSV replication in both starter cells and downstream target cells by establishing an antiviral state in distant brain regions (Drokhlyansky et al 2017, van den Pol et al 2014). This may explain why IFNAR1 knockout is required for long-distance spread from the dorsal striatum to the substantia nigra: unlike local inhibition at the injection site, IFNAR1 knockout abolishes type I interferon signaling in both the injected region and downstream target regions. Although the requirement for an interferon-deficient genetic background represents a substantial practical bottleneck, our findings indicate that broad suppression of type I interferon signaling is necessary for long-distance VSVdG spread (Figure 7).

**Figure 7.**
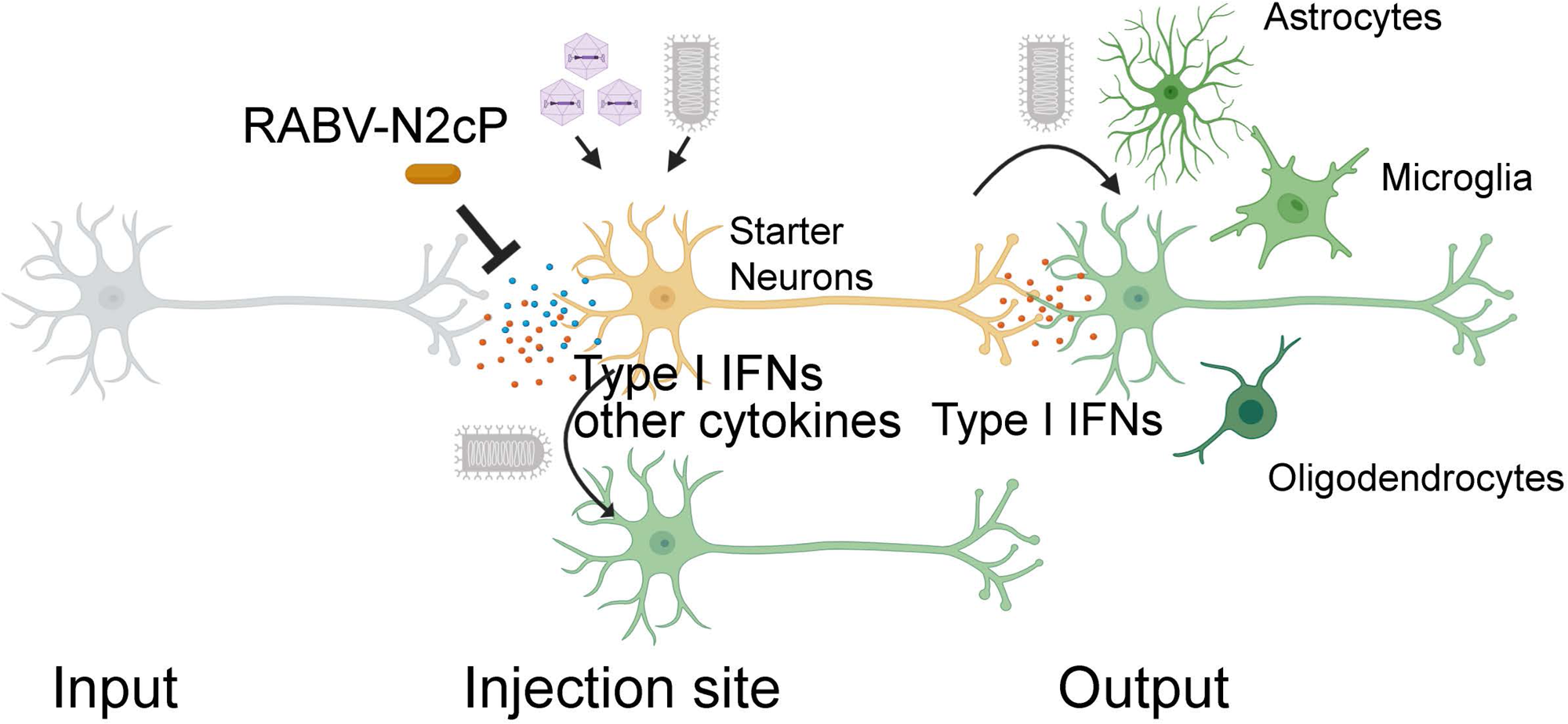
Proposed model for VSVdG-based one-step anterograde transmission VSVdG-based one-step anterograde spread requires VSV replication and VSV-G complementation in starter neurons but is restricted by antiviral cytokine responses at both the injection site and downstream output regions. At the injection site, AAV transduction and VSV infection may induce type I interferons and other cytokines that inhibit local VSV replication and spread. RABV-N2cP or a cytokine-blocking antibody cocktail can antagonize these antiviral barriers and permit VSVdG spread from starter neurons. In downstream projection targets, type I interferon signaling may further restrict infection of output-region cells. Therefore, long-distance spread was observed only in IFNAR1-knockout or IFNAR1/IFNGR1 DKO mice. However, because the VSV receptor LDLR is expressed by all major brain cell types, including neurons and glial cells, and has been reported to localize predominantly to the somatodendritic domain of neurons, VSVdG released from starter neurons can infect both neurons and glial cells in downstream regions but is unlikely to infect upstream input neurons, resulting in one-step anterograde spread.

Type I/type II interferon-independent cytokine signaling was also strongly implicated tin restricting VSVdG-Md spread. Three lines of evidence support this conclusion. First, RABV-N2cP was required even in IFNAR1/IFNGR1 DKO mice for efficient spread. Second, the different activities of the two RABV-P genes point to non-interferon effects. Only RABV-N2cP, but not RABV-B19P supported efficient VSV spread from dorsal striatum to substantia nigra. These differences cannot be ascribed to interferon antagonism, as both RABV-N2Cp and RABV-B19P inhibit nuclear translocation of activated STAT1, a key mediator of type I and type II interferon signaling (Brzozka et al 2006, Hossain et al 2019, Vidy et al 2005, Vidy et al 2007). Third, a cytokine-blocking antibody cocktail, targeting nine cytokines (not including anti-interferons), can substitute for AAV-RABV-N2cP. Further work will be needed to determine which cytokines are most important (Figure 7).

Microglia depletion showed context-dependent effects. With VSV-G–coated VSVdG, microglia depletion was required together with IFNAR1/IFNGR1 DKO and AAV-RABV-N2cP. In contrast, EnvA-pseudotyped VSVdG-Md spread efficiently without microglia depletion, suggesting that the difference arises during initial infection. Microglia express high levels of TLR4 and CD14 and are therefore well positioned to detect VSV-G on viral particles (Georgel et al 2007, Zhang et al 2014). Although virion-associated VSV-G has been reported to preferentially activate type I interferon signaling in macrophages (Georgel et al 2007), our results suggest that VSV-G–dependent activation of microglial TLR4 may also trigger release of other inflammatory cytokines. This possibility is supported by the observation that microglia depletion remained necessary even in IFNAR1/IFNGR1 DKO mice, implying that cytokines beyond type I and type II interferons contribute to the restriction of VSV spread in this context. Consistent with this interpretation, TLR4 activation in microglia has been shown to induce release of pro-inflammatory cytokines such as TNF-α, IL-1α, and IL-1β (Liddelow et al 2017, Zhou et al 2006).

An important strength of this system is that VSVdG-Md spread was detected in expected striatal output regions, including the GPe, GPi, and substantia nigra, but was not broadly observed in other brain regions. SNc labeling following striatal injection could, in principle, result from either anterograde or retrograde transmission. Striatal patch neurons project directly to the SNc, whereas SNc neurons project back to the striatum through the nigrostriatal dopaminergic pathway (Gerfen 1984, Matsuda et al 2009, Nelson & Kreitzer 2014, Watabe-Uchida et al 2012). Similarly, the striatum and GPe are reciprocally connected. Striatal projection neurons innervate the GPe, whereas a molecularly distinct population of arkypallidal GPe neurons provides dense GABAergic feedback to the dorsal striatum (Abdi et al 2015, Mallet et al 2012, Mallet et al 2016).

Several observations nevertheless argue against substantial retrograde transmission along long-range projections. Following striatal injection, we did not detect VSVdG-Md labeling in the cortex or thalamus, even though these regions are major sources of input to the striatum and are readily identified by monosynaptic retrograde tracing (Guo et al 2015, Wall et al 2013). Our results (Figure 1C) and previous studies have found that VSV-G–coated VSVdG produces only local neuronal cell-body labeling following intracranial injection (van den Pol et al 2009). Moreover, VSV-G–pseudotyped lentiviral vectors did not produce detectable neuronal cell-body labeling in distal projection regions when examined after survival periods ranging from approximately two weeks to several months, arguing against both rapid and substantially delayed retrograde transport (Baekelandt et al 2002, Humbel et al 2021, Mazarakis et al 2001). By contrast, RABV-G coated RABVdG and lentiviral vectors can retrograde label neurons from axonal terminals (Kato et al 2007, Mazarakis et al 2001, Wickersham et al 2007a). Neither our experiments nor the lentiviral studies, however, exclutde rare retrograde infection or retrograde labeling within local microcircuits. Neurons infected through nearby axon terminals would be difficult to distinguish from neurons infected directly at their somata near the injection site. Nevertheless, the absence of labeling in major long-range striatal input regions, together with robust labeling in established striatal output targets, supports the conclusion that VSVdG-Md spread was predominantly anterograde over long distances.

It is important to note several limitations of this VSV tracing system. First, VSV replication itself caused unhealthy morphology in starter cells, even in the absence of VSV-G expression. Second, VSV-G expression at axon terminals appears to induce local stress, as suggested by reduced NeuN immunoreactivity in downstream regions. Thus, although VSV-G fusogenic activity is required for viral spread, it may also perturb target regions even within a short experimental window. Such stress-induced reductions in cell-type markers may further complicate the identification and classification of labeled cells in viral tracing studies.

Another major limitation is that VSVdG-Md labeled both neurons and glial cells in downstream regions. Because monosynaptic tracing is generally intended to identify synaptically connected neurons (Callaway & Luo 2015), glial infection complicates interpretation. The presence of EGFP+ cells positive for NeuN, SOX9, SOX10, and Iba1 suggests that VSVdG-Md can infect multiple cell types near starter-cell axon terminals. VSV receptors have been identified as the LDL receptor (LDLR) and related family members (Finkelshtein et al 2013). In the brain, LDLR is expressed by all major cell types, including neurons, glial cells, and endothelial cells (Zhang et al 2014), suggesting that once virions are released from axon terminals and gain access to nearby cells, they may enter cells of multiple types. Whether such entry progresses to productive infection and results in detectable EGFP expression, however, is likely to depend on the antiviral state of the target cells, which is shaped by antiviral cytokine exposure. In this context, microglial labeling in the substantia nigra is consistent with our previous finding that productive VSVdG-Md infection of microglia was higher in IFNAR1/IFNGR1 DKO and IFNAR1-knockout mice, compared with wild-type and IFNGR1-knockout mice (Krause & Cepko 2024). These observations further support the idea that type I interferon signaling is a major determinant of VSV spread.

LDLR has been reported to localize predominantly to the somatodendritic domain of cultured neurons (Jareb & Banker 1998), and VSV-G has likewise been shown to cluster in dendritic regions in cultured neurons (Dotti & Simons 1990). Our finding that VSV-G supported both intrastriatal spread and long-distance spread to established projection targets suggests that, in vivo, AAV-driven overexpression of VSV-G may result in its localization not only to somatodendritic compartments but also to axons or presynaptic terminals. The strong anterograde bias of long-distance spread in our system may also be influenced by the subcellular distribution of LDLR-family receptors. Preferential receptor availability on somatodendritic membranes could facilitate infection of downstream recipient cells while limiting retrograde infection through the incoming axon terminals of neurons projecting to the injection site. Broad expression of LDLR-family receptors across neural cell types may additionally contribute to the eventual labeling of both neurons and glial cells in downstream regions (Figure 7). Nevertheless, cultured neurons may not fully reproduce the polarity and membrane-protein trafficking of neurons in vivo, and distinct neuronal populations may sort both VSV-G and LDLR-family receptors differently. Thus, the respective contributions of viral glycoprotein localization and receptor polarity to the direction and cellular specificity of VSVdG-Md spread remain open questions.

Overall, this study establishes that VSVdG can be adapted for one-step anterograde tracing in vivo when viral cytotoxicity and host antiviral immunity are carefully controlled. However, the system remains limited by its dependence on IFNAR1-knockout mice, residual cytotoxicity, and labeling of both neurons and glial cells in downstream regions。

Further optimization will therefore be required to reduce cytotoxicity, eliminate the requirement for an interferon-deficient genetic background, and improve neuronal specificity.

## Author Contributions

X.M. and C.L.C. designed research; X.M. performed research; X.M. and C.L.C. analyzed data; and X.M. and C.L.C. wrote the paper.

## Acknowledgments

We thank HHMI for funding this study. We are grateful to Christopher L. Parks at the International AIDS Vaccine Initiative (IAVI) for providing the plasmids used for VSV rescue, including those encoding VSV N, P, L, M, and G, as well as T7 polymerase. We also thank Dr. Paula Montero Llopis and Dr. Praju Vikas Anekal of the MICRON Imaging Core at Harvard Medical School for their expert advice on imaging.

