## Supplementary material for "Combined Disruption of Multiple Cytokine Signaling Pathways Enables One-Step Anterograde Tracing with Vesicular Stomatitis Virus": Figure S1, Figure S2 (from another version of this preprint: version 1, 2026-05-25)

### Supplementary Materials

.

Supplementary Figure 1

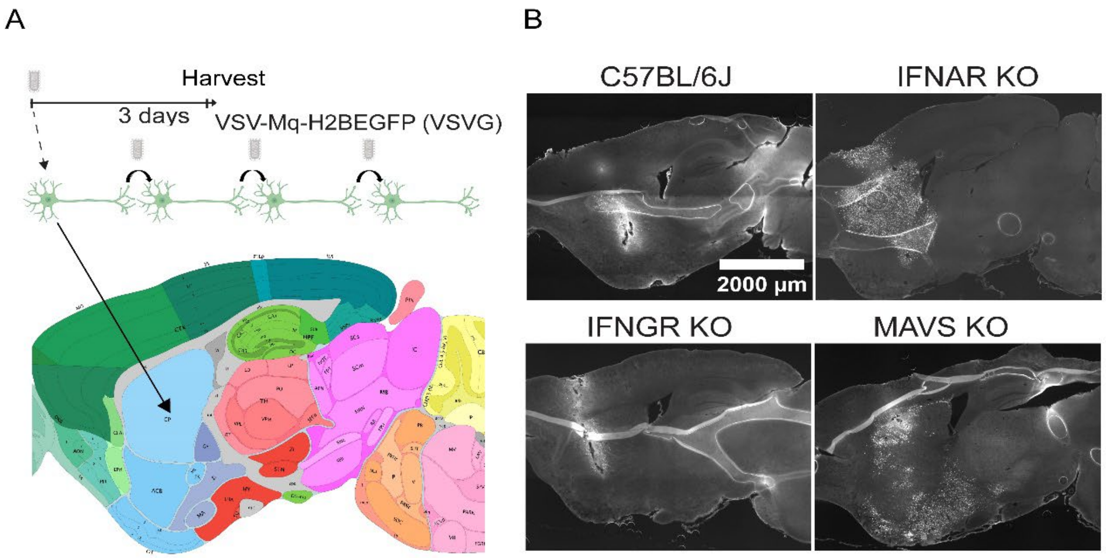


Figure S1. Multistep anterograde spread by VSV-Mq-H2B-EGFP (VSV-G) in wild-type and immune-deficient mice

(A) Experimental design.

(B) Representative images of sagittal section of the mouse brain showing H2B-EGFP labeling after VSV infection in the indicated mouse line. n=2 each.

Supplementary Figure 2

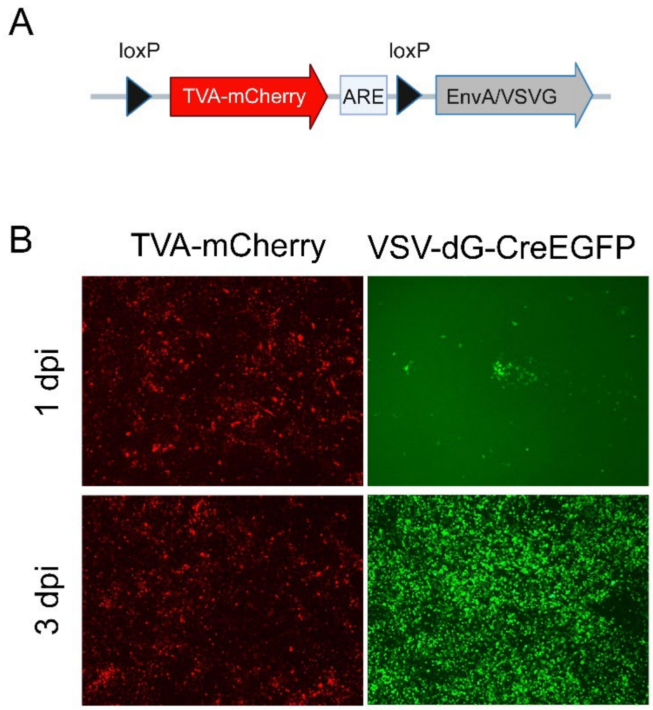


Figure S2. Generation of VSVdG-Md-CreEGFP(EnvA)

(A) Plasmid used to generate VSVdG-Md-CreEGFP(EnvA). After VSVdG-Md-CreEGFP enters the cell, Cre-mediated recombination removes the TVA-mCherry cassette, allowing EnvA expression alone. With the aid of this plasmid, both VSVdG-Md-CreEGFP(VSV-G) and VSVdG-Md-CreEGFP(EnvA) can be amplified from a low multiplicity of infection (MOI).

(B) Representative images showing VSVdG-Md-CreEGFP(EnvA)-infected cells forming a small colony at 1 day post-infection (dpi) and spreading throughout the dish by 3 dpi after infection at a MOI of 0.01.
